# Design, Synthesis, and Biological Evaluation of Novel Mitochondria-targeting Exosomes

**DOI:** 10.1101/2023.07.04.547719

**Authors:** Xin Yan, Xinqian Chen, Zhiying Shan, Lanrong Bi

## Abstract

Mitochondrial dysfunction is implicated in both brain tumors and neurodegenerative diseases, leading to various cellular abnormalities that can promote tumor growth and resistance to therapies, as well as impaired energy production and compromised neuronal function. Developing targeted therapies aimed at restoring mitochondrial function and improving overall cellular health could potentially be a promising approach to treating these conditions. Brain-derived exosomes (BR-EVs) have emerged as potential drug delivery vessels for neurological conditions. Herein, we report a new method for creating mitochondria-targeting exosomes and test its application in vitro and in vivo.

## Introduction

Brain diseases represent a highly intricate and multifaceted challenge for medical professionals. Conditions like brain tumors, stroke and neurodegenerative disorders are complex to manage, with the blood-brain barrier (BBB) acting as a formidable obstacle that can hinder targeted treatment of affected areas in the brain.^1^ The BBB can make it challenging to effectively target diseased cells, which can impede the efficacy of treatment.^2^ This challenge is particularly evident in the case of brain tumors, which can be highly invasive and difficult to remove.^3^ Early diagnosis of neurodegenerative diseases can also be problematic, leading to limited treatment options and reduced chances of successful intervention.^4^ Despite the challenges, there is hope for the future. Researchers are constantly exploring new ways to overcome the BBB, target diseased cells, and enhance therapeutic efficacy to unlock new treatment possibilities and improve patient outcomes.

As the field of brain tumors and neurodegenerative diseases progresses, one promising area of focus is the targeting of mitochondria. These powerhouses of the cell are responsible for producing energy and regulating cellular metabolism, making them a crucial component of overall cellular health. Dysfunctional mitochondria have been linked to a range of diseases, including Alzheimer’s (AD),^5^ Parkinson’s (PD), ^5^ and brain tumor. ^6^ There are several reasons why targeting mitochondria may be beneficial in the treatment of these conditions.^7,8^ For instance, enhancing ATP production can help restore energy balance within cells, which is often disrupted in neurodegenerative diseases.^9^ Additionally, reducing ROS (reactive oxygen species) production can protect cells from oxidative damage, which is thought to contribute to the development and progression of many diseases. Targeting mitochondria may also help regulate apoptosis (programmed cell death), which can be dysregulated in cancer cells.^10^ Protecting neurons from damage is another potential benefit of such therapies, as is repairing mitochondrial DNA mutations that can contribute to disease development. Furthermore, targeting mitochondria can enable more precise and efficient drug delivery, which may improve treatment outcomes and reduce side effects. While these potential benefits are promising, further research is necessary to develop effective and safe mitochondrial-targeted therapies.

In recent years, brain-derived exosomes (BR-EVs) have been gaining attention as a promising option for drug delivery in the field of neurology. These small extracellular vesicles possess the unique ability to naturally cross the BBB, allowing for tar-geted treatments to be administered to neurological conditions.^11^ Furthermore, BR-EVs are biocompatible and exhibit low immunogenicity, making them an ideal choice for drug delivery applications. ^11^ These features make BR-EVs an attractive therapeutic option for treating brain diseases. Our research team has hypothesized that by modifying the composition of the exosomal membrane, we can enhance targeting and improve drug loading efficiency, which could ultimately lead to more effective treatments for brain diseases. In this manuscript, we present our findings on the preparation and characterization of BR-EVs with mitochondria-targeting capabilities, which we believe could be a potential therapeutic strategy for brain tumors and neurodegenerative diseases.

## Results and Discussion

Preparing BR-EVs from rat brains is a complex process that involves several steps (**Fig. 1A**). Firstly, the rats are euthanized with an overdose of isoflurane, and their brains are carefully removed and cut into small pieces. The exosomes are then separated from other cellular components using ultracentrifugation. Finally, various techniques such as transmission electron microscopy (TEM), dynamic light scattering (DLS), MALDI-TOF mass spectrometry are employed to confirm the identity of the isolated exosomes.

**Figure 1.**
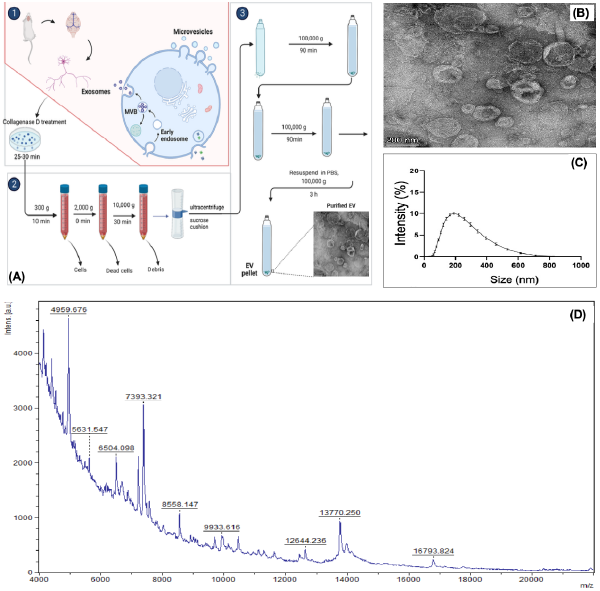
Preparation and characterization of the brain-derived exosomes, BR-EVs. (**A**) Preparation of modified exosomes (Rho-EVs) by ultracentrifugation; (**B**) TEM image of BR-EVs; (**C**) DLS analysis of BR-EVs; (**D**) The proteomic composition of exosomes was analyzed by MALDI-TOF mass spectrometry.

The size of unmodified and modified BR-EVs can affect their cellular uptake, stability, and functional properties, making investigations of their sizes distribution crucial. DLS is a relatively quick and non-destructive technique, making it suitable for measuring exosomes in their native state. It measures the fluctuations in dispersed light intensity resulting from the Brownian motion of particles in solution. ^12^

As shown in **Fig. 1C**, the hydrodynamic sizes of BR-EVs approximately are 120-200 nm. It’s important to keep in mind that the hydrodynamic sizes of exosomes measured by DLS include the size of the exosome itself as well as the surrounding water molecules and any solutes or biomolecules present in the suspension. This means that the measured size may not reflect the actual size of the exosome core alone. DLS provides information about the size distribution of exosomes in a sample. Therefore, additional complementary techniques, such as TEM is often used in conjunction with DLS to gain a more comprehensive understanding of exosomes’ characteristics.

As shown in **Fig. 1C**, BR-EVs typically range in size from approximately 30 to 150 nm. BR-EVs generally have a cup-shaped or saucer-like morphology, formed by the inward budding of the endosomal membrane (**Fig. 1B**). The characteristics of BR-EVs may vary in different biological contexts due to various factors such as cell type, cellular state, and environmental conditions. ^13^

MALDI-TOF mass spectrometry is a powerful tool for exosome characterization and has contributed to our understanding of exosome biology and their potential diagnostic and therapeutic applications. ^14^ As shown in **Fig. 1D**., the obtained mass spectra correspond to the proteins and other biomolecules present in the exosomes. Currently, we are analyzing the mass spectra and hope to gain insights into the protein composition and heterogeneity of exosomes, identify potential biomarkers, and compare exosome samples from different conditions or disease states, and detailed results will be reported elsewhere.

BR-EVs are composed of a lipid bilayer membrane that encapsulates various molecules, including proteins, nucleic acids, lipids, and metabolites.^11^ These molecules include some proteins that have amino groups on the exosomal surface, which can be modified chemically for various purposes. By modifying the BR-EVs chemically, we can enhance their functionality and customize them for specific applications. The modifications can be used to attach diagnostics or therapeutic agents to the exosomal surface, which allows for better targeting specificity, enables targeted delivery to specific organelles, and facilitates the visualization and tracking of exosomes in brain mito-chondria.

To improve their targeting specificity and therapeutic potential, we modified the amino groups on the surface of exosomes. More specifically, 4-pentynoic acid, N-hydroxy succinimide (NHS), and 1-Ethyl-3-(3-dimethylaminopropyl) carbodiimide (EDC) were used to introduce alkyne functionality to the surface of BR-EVs. The process involves activating the carboxylic acid groups on the BR-EVs’ surface with NHS and EDC, then adding activated 4-pentynoic acid to the solution to form stable amide bonds with primary amines on the BR-EVs’ surface. This results in covalently attached alkyne groups on the BR-EVs’ surface, which can be used for subsequent bioconjugation reactions. This approach allows for modifications or specific interactions with other molecules through a “click” chemistry. We utilized the alkyne functionality as a site for chemical conjugations with a mitochondrial-targeting ligand (MTL), rhodamine derivative **3** (**Fig. 2A**), via a “click” chemistry based on copper(I)-catalyzed azide-alkyne cycloaddition (CuAAC).^15,16^ Rhodamine derivatives have a unique structure with a central xanthene ring system that can be modified to add specific properties or targeting capabilities. Regarding mitochondrial targeting, certain rhodamine derivatives, have been developed to selectively accumulate in the mitochondria. This makes them valuable tools for studying mitochondrial function and dynamics. Finally, we isolated and purified the modified exosomes, Rho-BR-EVs, using a Sepharose CL-4B column. This versatile method of modifying exosomes using CuAAC has allowed us to create a new diagnosis and therapeutic tool.

**Figure 2.**
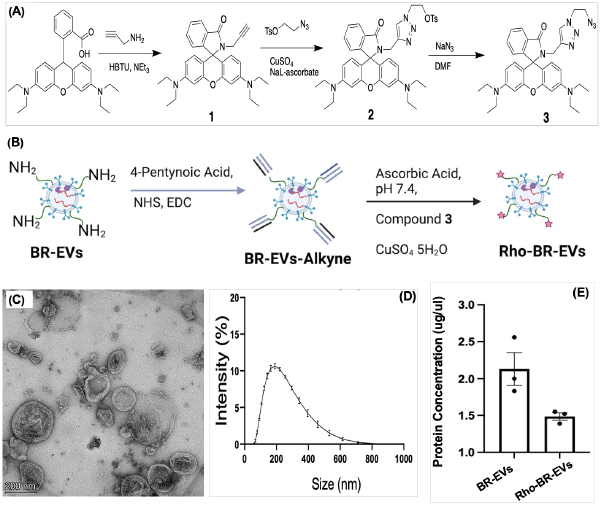
Preparation and Characterization of MTEs (Rho-BR-EVs) based on brain-derived exosomes. (**A**) Preparation of a mitochondria-targeting ligand (MTL), i.e., rhodamine-derived **3**; (**B**) Chemical modification of exosomal surface proteins using a copper(I)-catalyzed azide-alkyne cycloaddition (CuAAC). (**C**)TEM image of Rho-BR-EVs; (**D**) DLS analysis of Rho-BR-EVs; (**E**) Protein content analysis of Rho-BR-EVs and un-modified BR-EVs.

Upon conducting thorough TEM and DLS analyses both before and after the chemical modification process, we have found that the resulting modifications have not had a significant impact on the morphology (**Fig. 2C**) and size (**Fig. 2D**) of the exosomes. This leads us to confidently conclude that the chemical modification process is safe for use with regards to its effects on the physical characteristics of the exosomes. However, it is worth noting that these analyses only provide information on the size and morphology of the exo-somes. While these physical characteristics are certainly important, there may be other aspects of exosomes that require further evaluation using additional methods.

When working with modified exosomes, it’s important to examine the proteins present in them to fully understand the changes that occur during the modification process and evaluate their safety and efficacy for different applications. Through our analysis using Bradford protein assay, we observed some extent of protein loss (**Fig. 2E**), which is normal during the isolation and purification process. However, this may pose a problem during modification and purification. To minimize protein loss, it’s crucial to optimize the protocols and procedures used, and implement quality control measures to ensure accurate characterization of modified exosomes.

Next, we examined the subcellular localization of modified exosomes through the preparation and cultivation of primary neuron cells. To achieve this, we isolated primary neurons from brains of neonatal Sprague Dawley (SD) rats and cultured them in specialized neuronal culture media under controlled incubator conditions until they had matured. Upon maturity, we introduced Rho-BR-EVs into the culture media and allowed the cells to take up these modified exosomes.

Using confocal microscopy, we were able to obtain high-resolution images of cells to further investigate subcellular localization. To enhance visualization, we counterstained the cultures with MitoTracker, a fluorescent mitochondria-specific marker. Upon merging the confocal images of Rho-BR-EVs and MitoTracker, we were able to successfully observe co-localization with the MitoTracker, which indicated that the modified exo-somes had targeted the intended subcellular location. To gain insights into the behavior of Rho-BR-EVs in cells, we conducted a time course study over 72 hours, closely monitoring changes in fluorescence intensity and localization (**Fig. 3**). It appears that the uptake of Rho-BR-EVs in primary neuron cells is dependent on time, and the fluorescence intensity increases over time. This finding suggests that the uptake of exosomes may indeed increase over an extended period of time. Exo-somes are important for communication between cells as they can transfer various molecules such as proteins, nucleic acids, and lipids. In our present study, we observed the fluorescence intensity of Rho-BR-EVs increased with time, indicating anaccumulation of these modified exosomes within the cells or an increasing rate of uptake as time progressed. Upon analyzing these results, we hypothesize that endocytosis was likely the primary method of Rho-BR-EV uptake by cells. Once internalized through endocytosis, the Rho-BR-EVs must be able to escape the endosomal compartment and target the mitochondria, as shown in **Fig. 4**.

**Figure 3.**
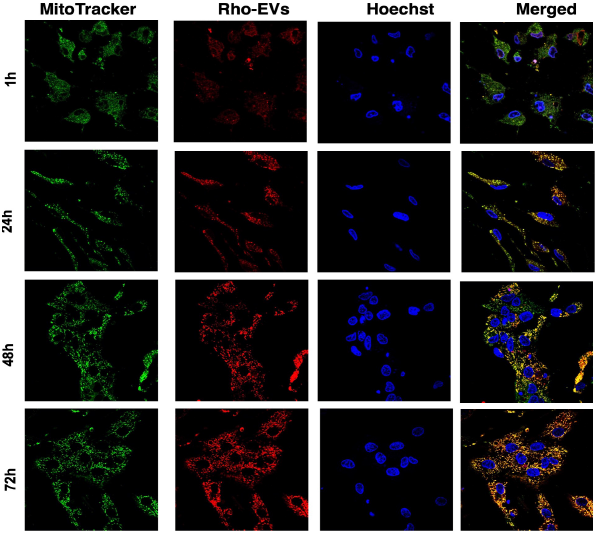
Representative confocal fluorescent pictures of primary neurons incubated with Rho-BR-EVs (2 μg/mL, red fluorescence), MitoTracker (80 nM, green fluorescence), Hoechst 33242 (0.1 μL/mL, blue), and merged images (yellow). The Rho-BR-EV time-course investigation was carried out at various time points (1h, 24h, 48h, and 72h). The fluorescent pictures were captured using a confocal laser scanning fluorescence microscope with a 60 × objective lens and medium free of FBS and phenol-red.

**Figure 4.**
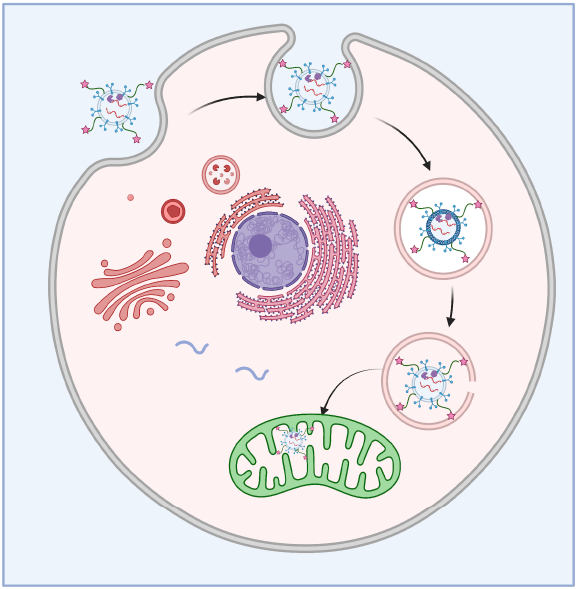
It is assumed that cells use endocytosis to absorb Rho-BR-EVs. After entering the cell, the Rho-BR-EVs must break free from the endosomal compartment and target the mitochondria. Rho-BR-EVs tend to group in a specific manner within the mitochondria, likely due to the unique properties of the rhodamine derivative, which can pass through the mitochondrial membrane and is attracted to the negative charge on the inner membrane, resulting in the characteristic punctate pattern of Rho-BR-EVs within the mitochondria.

It’s important to keep in mind that brain tumors and neurodegenerative diseases can have a significant impact on the hippocampus and cortex regions of the brain. These regions play a crucial role in higher cognitive functions, like memory formation, spatial navigation, language, attention, and executive functions. Unfortunately, these conditions can disrupt these functions and cause impairments. For instance, brain tumors can directly affect the hippocampus and cortex, while neurodegenerative diseases can indirectly impact them through abnormal protein buildup. That’s why we carefully studied the uptake and distribution of Rho-BR-EVs in these regions.

Modified brain exosomes were delivered into rat brain through intracerebroventricular (ICV) injection. 24 hours following ICV injection, rats were euthanized, their brains were sectioned, and exosomes distribution were observed as we detailed in the Materials and Methods section. Our findings showed that Rho-BR-EVs were present in the hippocampus (**Fig. 5**) and cortex (**Fig. 6**), indicating that cells like neurons and glial cells had absorbed them.

**Figure 5.**
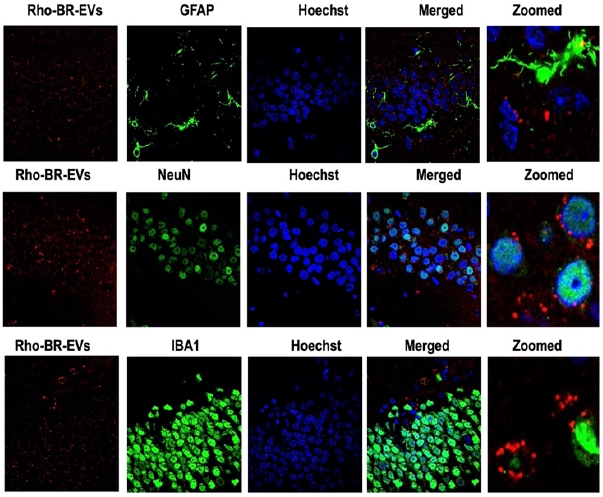
Rho-BR-EV visualization in hippocampal regions: Images of confocal laser scanning of rat brain slices fixed with 4% PFA. The red channels indicate fluorescently labeled exosomes (Rho-BR-EVs) throughout the picture, whereas the green channels reflect GFAP-(row I), NeuN-(row II), and IBA1-positive (row III) cells in brain slices, respectively. On the right, zoomed-in images are exhibited. The fluorescent images were captured using a 100 × objective lens on a confocal laser scanning fluorescence microscope.

**Figure 6.**
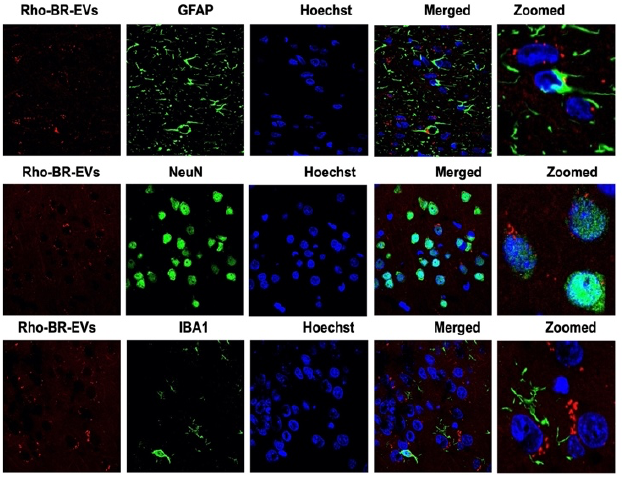
Rho-BR-EV visualization in cortical regions: Images of confocal laser scanning of rat brain slices fixed with 4% PFA. The red channels indicate fluorescently labeled exosomes (Rho-BR-EVs) throughout the figure, whereas the green channels reflect GFAP-(row I), NeuN-(row II), and IBA1-positive (row III) cells in brain slices, respectively. On the right, zoomed-in images are exhibited. The fluorescent pictures were captured using a 100 × objective confocal laser scanning fluorescent microscope.

Next, we utilized primary antibodies against NeuN, a neuron-specific nuclear protein, Ionized calcium binding adaptor molecule 1 (IBA1), a microglia-specific calcium-binding protein, and glial fibrillary acidic protein (GFAP), a marker for astrocytes to track the cell types absorbing Rho-BR-EVs. It was fascinating to note that Rho-BR-EVs were taken up by all three types of cells, including neurons, astrocytes, and microglia in hippocampus (**Fig. 5**) and cortex (**Fig. 6**). However, administering Rho-BR-EVs did not lead to any significant alterations in the expression levels of GFAP or IBA1 in the hippocampus or cortex regions compared to the control groups with vehicles. These results indicate that Rho-BR-EVs could be a potential drug carrier for treating brain damage without causing significant changes in glial cell activity.

## Conclusion

We have developed a new method for preparing exosomes that can specifically target mitochondria, and we have tested this approach both in vitro and in vivo. This exciting breakthrough has significant potential for improving targeted drug delivery by enabling us to deliver drugs directly to specific tissues, cells, and organelles within the body. Furthermore, our discovery of Rho-BR-EVs within mitochondria could have far-reaching implications for studying mitochondrial function or dysfunction. By investigating the role of mitochondria-targeting exosomes in intercellular communication and signaling within the brain, we may be able to develop a new therapeutic strategy for brain tumor and neurodegenerative diseases.

Our research has successfully demonstrated the subcellular localization of modified exosomes in primary neuron cells, highlighting the immense potential of these modified exosomes for targeted drug delivery purposes. This approach could revolutionize the field of drug delivery, enabling the targeted delivery of drugs to specific subcellular locations, thereby enhancing therapeutic efficacy while minimizing adverse effects. While the therapeutic potential of mitochondrial targeting exo-somes is still being investigated, these vesicles offer promise as a novel and tailored strategy for treating brain tumor and neurodegenerative diseases.

## Limitation

At the beginning of our project, we assumed it challenging to determine the targeting specificity and fluorescent intensity of modified exosomes in brain tissues due to the complexity of brain tissue and high auto-fluorescence of brain regions. To address this issue, we initially administered Rho-BR-EVs directly into the cerebrospinal fluid (CSF) within the brain’s ventricles using ICV injection. Our initial data suggest that Rho-BR-EVs could travel through the CSF circulation to different brain regions and be taken up by various types of cells, including neurons, astrocytes, and microglia. This indicates the potential role of Rho-BR-EVs in delivering drugs to the brain. However, it is important to note that ICV injection used in our present study is an invasive procedure that requires specialized equipment and skills for safe execution. Furthermore, it may pose risks of infection, inflammation, and damage to brain tissue. Nevertheless, we have recently demonstrated that Rho-BR-EVs can cross the BBB through peripheral administration, such as intravenous or intraperitoneal injection (data not shown). This promising and safe new drug delivery system to the central nervous system can be used to treat brain diseases. We will report our new findings elsewhere.

## Materials and Methods

### Procedure for synthesizing compound 1: ^17^

In 20 ml of anhydrous dichloromethane, 0.90g (1.80 mmol) of Rhodamine B was dissolved. Propargylamine (0.12ml, 2.0mmol), 2-(1H-Benzotriazol-1-yl)-1, 3, 3-tetramethyluronium-hexafluoro-phosphate (HBTU) (0.783g, 2.0mmol), and triethylamine (1ml, 5.76mmol) were added to the mixture. The mixture was stirred at room temperature overnight. The mixture was rinsed three times with brine after being diluted with 20ml dichloromethane. The organic layer was then vacuum evaporated, filtered, and dried over anhydrous sodium sulfate. The final product was purified by flash column chromatography on silica gel (Hexane: EtOAc, 4:1) to produce compound 1 (0.86g, 80%) as a light pink solid. ^1^H NMR (400 MHz, CDCl_3_) δ ppm 1.14 (t, *J* = 7.2 Hz, 12 H), 1.74 (t, *J* = 2.4 Hz, 1H), 3.32 (q, *J* = 7.2 Hz, 8H), 3.93 (d, *J* = 2.4 Hz, 2H), 6.25 (dd, *J* = 8.8, 2.8 Hz 2H), 6.37 (d, J =2.8 Hz, 2H), 6.45 (d, *J* = 8.8 Hz, 2H), 7.10 (m, 1H), 7.41 (m, 2H), 7.91 (m, 1H). ^13^C NMR (101 MHz, CDCl_3_) δ 189.81, 167.47, 153.89, 153.61, 148.99, 132.78, 130.60, 129.27, 128.16, 123.95, 123.18, 108.22, 105.33, 98.03, 78.54, 70.27, 65.04, 44.68, 28.84, 12.91.

### Synthetic procedure for compound **2**: ^17^

In 25ml of THF, mix together compound **1** (0.5g, 1.00 mmol) and 2-azidoethyl 4-methylbenzenesulfonate (0.45 g, 1.25 mmol): Copper (II) sulfate (0.21 mmol, 40 mg) and sodium ascorbate (0.42 mmol, 82 mg) were added to tert-butanol:water = 3:1:1. After that, the reaction mixture was stirred at room temperature for 6 hours. The reaction was rinsed three times with 10ml saturated NaHCO_3_ before being extracted with 40ml ethyl acetate. The organic layers were mixed, dried over anhydrous sodium sulfate, filtered, and concentrated. The crude residue was purified by flash column chromatography on silica gel (eluent: Hexane: EtOAc = 1:4) to get compound 2 (0.62g, 80%) as colorless foam.^1^H NMR (400 MHz, CDCl_3_) δ ppm 1.09 (t, *J* = 7.0 Hz, 12H), 2.32 (s, 3H), 3.26 (q, *J* = 7.0Hz, 8H), 4.21 (t, *J* = 5.0 Hz, 2H), 4.29 (t, *J* = 5.0 Hz, 2H), 4.37 (s, 1H), 6.12 (dd, *J* = 8.8, 2.0 Hz,2H), 6.24 (d, *J* = 8.8 Hz, 2H), 6.31 (d, *J* = 2.0 Hz, 2H), 6.84 (s, 1H), 7.10-6.94 (m, 1H), 7.21 (d, *J* = 8.5 Hz, 2H), 7.47-7.29 (m, 2H), 7.65-7.46 (m, 2H), 8.09-7.73 (m, 1H). ^13^C NMR (100 MHz, CDCl_3_) δ ppm 168.03, 153.04, 153.61, 148.95, 145.64, 144.76, 132.79, 132.23, 131.19, 130.26, 129.03, 128.32, 127.98, 124.14, 123.40, 123.09, 108.13, 105.50, 98.01, 76.92, 67.48. 65.19, 48.60, 44.53, 35.58, 21.84, 12.81.

### Synthetic procedure for compound **3**:^17^

NaN_3_ (70 milligram, 1.00 mmol) was added to a solution of compound 2 (0.50 g, 0.70 mmol) in 10 ml DMF while stirring. Overnight, the reaction mixture was gently re-fluxed. The reaction mixture was rinsed three times with 10 ml of saturate NaHCO_3_ and extracted with 40 ml of EtOAc. The organic layers were combined, desiccated over anhydrous Na_2_SO_4_, filtered, and evaporated in a vacuum. Compound **3** (0.40g, 96%) was obtained as a white froth after the crude residue was purified by flash column chromatography on silica gel (eluent: Hexane: EtOAc=1:3).^1^H NMR (400 MHz, CDCl_3_) δ ppm 1.14 (t, *J* = 7.2 Hz, 12H), 3.30 (q, *J* = 7.2 Hz, 8H), 3.60 (t, *J* = 6.1 Hz, 2H), 4.20 (t, *J* = 6.1 Hz, 2H), 4.47 (s, 2H), 6.15 (dd, *J* = 8.9, 2.5 Hz, 2H), 6.30-6.33 (m, 4H), 6.95-7.18 (m, 3H), 7.37 (m, 2H), 7.86 (m, 1H). ^13^C NMR (101 MHz, CDCl_3_) δ 167.94, 153.56, 153.54, 148.82, 144.68, 132.71, 131.08, 128.96, 128.24, 124.03, 123.19, 123.02, 108.00, 105.54, 98.02, 65.25, 60.60, 50.68, 49.00, 44.60, 35.54, 19.48, 14.53, 12.94.

### Animals

Adult SD rats used in this study were purchased from Charles River Laboratories (Wilmington, MA). All animals were obtained on a 12:12-h light-dark cycle in a climate-controlled room, and chow and water were provided *ad libitum*. This study was carried out in strict accordance with the recommendations in the Guide for the Care and Use of Laboratory Animals of the National Institutes of Health. The protocol was approved by the Michigan Technological University Institutional Animal Care and Use Committee.

#### Preparation of brain-derived exosomes (BR-EVs)

Extracellular vesicles (EVs) were isolated based on established protocols with some modifications. ^19, 20^ Fresh rat brains (whole brains without cerebellum) preserved in PBS were cut into minute pieces on ice. Following the transfer, tissue fragments weighing 200 mg were placed in a 6-well plate containing 2 ml of Hibernate E (Gibco) in each well and were carefully dissociated until all the pieces were the same size: approximately 2 mm on one side, 2 mm on the length, and 2 mm on the height. Subsequently, 40 μl of Collagenase D (Sigma-Aldrich) and 4 μl of DNase I (Sigma-Aldrich) were added to each well to reach a final concentration of 2 mg/ml for Collagenase D and 40 U/mL for DNase I. The plate was immediately transferred into a pre-warmed 37 °C incubator under mild agitation (70 rpm) for 30 minutes. Immediately following the incubation step, the plate was placed back on the ice, and protease and phosphatase inhibitors (Sigma-Aldrich) were added to each well. Tissue fragments containing Hibernate E drained into a 50 ml tube by gravity through a 70 μm sterile cell strainer. Additional 1-2 ml PBS was added to each well to transfer the remaining tissue fragments. Differential centrifugation was performed on the filtered samples: I. 300 × g, 10 minutes, 4 °C; II. 2000 × g, 20 minutes, 4 °C; and III. 10,000 × g, 30 minutes, 4 °C. To remove larger particulates from the sample, the supernatants were collected and filtered through a 0.22 μm filter (Grainger Industrial Supply). The filtered supernatants were progressively layered over 4 ml of a 30% sucrose solution prepared in PBS before being centrifuged at 100,000 × g for 90 minutes at a temperature of 4 °C. (Optima XL-90 ultracentrifuge, SW 28 centrifuge rotor, Beckmann-Coulter). The supernatant was discarded, and the sucrose layer (approximately 5 ml) and pellets were collected. Additional ultracentrifuge (100,000 g, 90 minutes, 4°C) was performed and the pellets were resuspended in 50 μl of ice-cold PBS for subsequent steps. Total proteins of BR-EVs were extracted with RIPA buffer containing 0.5% PMSF (Sigma Aldrich). In brief, BR-EVs were subjected to an equal volume of supplemented RIPA buffer and sonicated 3 times each 5 seconds with each 5 second interval. Sonicated BR-EVs samples were lysed on ice for 20 minutes and mixed with pipetting every 5 minutes. Protein quantification was determined by Bradford reagent (Sigma Aldrich).

#### Preparation and culture of primary neuron cells

In neuroscience research, primary neuron cells derived from postnatal rat brains are extensively used as an in vitro model to explore neuronal development, function, and various neurological disorders. ^25,26^

Neurons were isolated from brains of 1-day-old SD pups using papain dissociation system (Worthington Bio-chemical Corporation) following manufacturer’s instruction. First, SD pups were euthanized using overdose of isoflurane, and their whole brains were removed and placed in a sterile culture dish containing cold PBS. The blood vessels and meninges that surround the brain were removed, the cerebellum and brainstem were taken away. The remaining brain region containing the neurons of interest (e.g., hippocampus, cortex) were transferred to a new culture dish, and cut into small (1-2 mm) pieces. In a sterile tube containing pre-warmed papain solution, place the tissue pieces and incubate the tube containing the tissue at 37°C with constant agitation for 60 minutes. After this enzymatic reaction, the mixture was gently triturated using a sterile Pasteur pipette to release cells. The cloudy cell suspension was centrifuged to collected cells. The cells were further purified with discontinuous density gradient centrifugation as detailed in the manufacturer’ manual. Cell concentration was then determined using cell counting equipment. The cell suspension was then diluted with neurobasal media supplemented with B27 serum to reach the desired cell density. The cells were then plated onto poly-D-lysine-coated culture dishes at the appropriate cell densities and volumes and incubated in 5% CO_2_ incubator for 7-10 days and ready for experiment. Half of the culture medium were replaced with fresh medium every 3 days. The cells’ growth and well-being are monitored on a regular basis using a microscope.

#### Modification of exosomes’ surface using NHS ester chemistry

Alkyne-modified exosomes: After adding 290mg propargylacetic acid (3 mmol) and 350mg n-hydroxysuccinimide (3 mmol) to 10 ml of PBS with stirring, the resulting solution was adjusted to pH 7.4 with sodium bicarbonate. After 1 hour of stirring in an ice bath, 460 mg of 1-ethyl-3-(3-carbodiimide dimethyl aminopropyl) was added, followed by another hour of stirring in an ice bath. Then, 100 μl of the reaction mixture was added to 250 μl of the exosome solutions, followed by 24 hours of shaking at room temperature. Exosomes were isolated from excess reaction material using a pH 7.4 PBS-conditioned Sepharose CL-4B column.

#### Preparation of Rho-BR-EVs via a “click” chemistry based on copper(I)-catalyzed azide-alkyne cycloaddition (CuAAC)

Labeling alkyne-modified exosomes with azido-rhodamine (compound 3) is a two-step process. First, the exosomes are labeled with compound 3 via a copper-catalyzed “click” chemistry reaction, and then the labeled exosomes are purified to remove any unbound dye and excess reactants.

The alkyne-modified exosomes are thawed and diluted to a final concentration of 1-2 mg/mL in PBS buffer. In a microcentrifuge tube, combine the following components to make the copper-catalyzed click chemistry reaction mixture: azido-rhodamine (compound 3), 10-20 μg per mL of exosome solution. Copper sulfate, 50 μM; Sodium ascorbate, 100 μM; TCEP, 1 mM; DMSO, 10% final concentration; Briefly agitate the reaction mixture, then add it to the exosome solution while stirring gently. Protected from light, incubate the reaction mixture at room temperature for 1-2 hours. After incubation, add 10 to 15 mL of PBS buffer to the reaction mixture and transfer it to an ultracentrifuge tube. To extract Rho-BR-EVs, centrifuge the reaction mixture at 100,000 × g for one hour at 4 °C. Resuspend the exosome particle in 1 mL of PBS buffer and discard the supernatant. Repeat the ultracentrifugation step to remove the unbound dye and superfluous reactants from the exosomes and to wash them. Reconstitute the exosome particle in 50-100 μL of PBS buffer and discard the supernatant.

#### Dynamic light scattering (DLS) analysis of exo-somes

Light scattering in resuspended pellets was diluted with DPBS to achieve a final protein concentration of 0.01 μg/μl. Diluted samples was put into a low-volume cuvette, and the size distribution of exo-somes was determined using the Malvern Zetasizer nano series. Cuvettes with lids were mixed upside down to avoid large particles from remaining at the bottom. In each sample, the measurement was repeated three times.

#### Morphology analysis by TEM

30 μl resuspended brain-derived sEVs were mixed with an equal volume of 2% paraformaldehyde (PFA) for 5 minutes. 5 μl of specimen was loaded on the carbon side of the grid and left for 1 min. Subsequently, the sample was gently blotted dry with a clean filter paper. Negative staining was performed with uranyl acetate (UA). A 3 μl fresh 2% UA solution was placed on the grid, left for 30 s and excess UA solution was blotted with another clean filter paper. Air-dried samples were visualized with FEI 200kV Titan Themis STEM.

#### MALDI-TOF mass spectrometry analysis

To carry out the MALDI-TOF mass spectrometry analysis, sinapinic acid (SA, 3,5-Dimethoxy-4-hydroxy-cinnamic acid) was used as a matrix to absorb the laser energy and facilitates the ionization of the analyte molecules. The exosome sample was mixed with the matrix solution, and a small aliquot of the mixture is deposited onto a MALDI target plate. The sample was then allowed to dry, forming a thin crystalline layer of the matrix with embedded exosome analytes. The target plate was then placed inside the MALDI-TOF mass spectrometer. A laser beam focused on the sample spot, causing the matrix to absorb the laser energy and desorb/ionize the exosome analytes. This generates positively charged ions from the exosome molecules. The generated ions were then separated based on their mass-to-charge ratio (m/z) as they travel through a flight tube.

#### Intracerebroventricular injection (ICV injection) of modified exosomes (Rho-BR-EVs)

Intracerebroventricular (ICV) injection is a method for administering medications or other substances directly into the brain’s cerebrospinal fluid (CSF) and allow them to circulate in the whole brain. ICV injection was performed as described in our previous publication.^27^ In brief, SD rats were anesthetized with 5% isoflurane and maintained at 2.5% throughout the surgery. Rats were placed stereotaxically over the heating device, and their rectal temperature was maintained at approximately 37 °C. The cranium was exposed by incising the scalp over the sagittal suture between the eyes and ears and removing the periosteum. The stereotaxic apparatus was adjusted to align bregma and lambda. A right hole was drilled through the cranium at the coordinates of the lateral ventricles.

The lateral ventricles had the following coordinates: 0.8 mm caudal to bregma, 1.6 mm in the mediolateral axis, and -3.6 mm in the dorsoventral axis. Using an UltraMicroPump 3 (World Precision Instruments), 5.5 μg of Rho-BR-EVs were injected into the lateral ventricle at a 1 μl/min rate. 24 hours after ICV administration, rats were perfused with ice-cold PBS and 4% paraformaldehyde (PFA). The rat brains were extracted, fixated overnight in a 4% PFA solution, and subjected to 30% sucrose in filtered PBS until the tissue sank. The brains were then embedded in O.C.T. compound (Sakura Finetek) and cryo-sectioned into 15 μm thick coronal sections for fluorescent immunostaining as described in the below.

#### Immunofluorescence staining

Immunostaining was performed as described in our previous publication ^28^ with some modification.. Brain sections containing the cortex or hippocampus were first washed in PBS three times for 10 minutes each, and then were incubated with 5% horse serum for 30 minutes. The sections were then incubated with either rabbit anti-NeuN antibody (Cell signaling, 1:300 dilution) or rabbit anti-GFAP antibody (Abcam, 1:500 dilution), or rabbit anti-IBA1 antibody (Fujifilm Wako, 1:500 dilution) in PBS containing 0.5% Triton X-100 and 5% horse serum for 48 h at 4°C. Afterward, brain sections were washed with PBS three times for 10 min each. Brain sections were then incubated with secondary antibody Alexa fluor 488 donkey anti-rabbit IgG at room temperature for 1h. The immunoreactivity of NeuN, GFAP, IBA1 as well as the fluorescence of Rho-BR-EVs were observed under a confocal microscope and images were taken.

